# Applying distinct CDMS strategies to observe non-classical virus capsid assembly

**DOI:** 10.64898/2026.04.29.721378

**Authors:** Lars Thiede, Anisha Haris, Tomislav Damjanovic, Jocky C.K. Kung, Jürgen Müller-Guhl, Ronja Pogan, Jennifer Rothe, Wiebke Schultze, Sanne Ugelstad, David Eatough, Kevin Giles, Steve Preece, Keith Richardson, Jakub Ujma, Charlotte Uetrecht

## Abstract

In conventional native mass spectrometry (MS), one faces severe limitations when challenged with heterogenous, high mass samples, commonly failing to resolve clear peak distributions and thus mass determination. Charge detection MS (CDMS) has emerged as a premier method to analyze these samples by determining mass-to-charge ratio (*m*/*z*) and charge (*z*) simultaneously. Here, the two currently available commercialized CDMS systems, the Orbitrap™-based Direct Mass Technology (DMT™) and the electrostatic linear ion trap (ELIT)-based Xevo™ CDMS are applied to human norovirus capsids from two different strains, GI.1 Norwalk and GII.17 Kawasaki. The norovirus capsid is highly heterogenous due to N-terminal processing on the repeating subunits that it is built from and commonly forms *T =* 3 and sometimes *T* = 4 particles. Both CDMS approaches were able to determine similar masses in both strains. GII.17 Kawasaki exhibits both *T* = 3 and *T* = 4 particles, though the Xevo CDMS measurements were closer to the theoretical mass than the DMT instrument. Interestingly, GII.17 Kawasaki also displayed non-classical mass distributions with high abundance in-between *T* = 3 and *T* = 4 which was then confirmed by cryogenic electron microscopy (cryo-EM), demonstrating an oval capsid shape. GI.1 Norwalk displays a wide mass distribution in both instruments that exceeds the theoretical *T* = 3 mass by 8-10 %. Proteomics and native MS experiments suggest possible interactions with a protein from the expression system. This study demonstrates the capabilities of two distinct CDMS methodologies on two viral capsids and presents the first non-classical capsid assembly in a GII.17 noroviral capsid.

## Introduction

Native mass spectrometry (nMS) is a branch of mass spectrometry (MS) that allows the analysis of intact biomolecular assemblies. The mass determination of macromolecular complexes in their native-like folded state can give detailed insights on their stoichiometries, composition and the dynamics of both^1–3^. This usually necessitates charge state resolution. Within a well-resolved mass-to-charge (*m*/*z*) spectrum, the charge state can be determined from the isotopic peak distribution or deconvolved from the charge state distribution, though this becomes difficult once large, heterogenous samples are introduced^4^. Mass heterogeneity can originate from a plethora of factors and is often found in large assemblies, like viruses and virus capsids^5^. Large assemblies have large surface areas that provide a high amount of interaction sites for adducts, for example metal cations or solvent molecules. High mass ions typically present at a higher *m/z*, limiting energy uptake in traditional collision induced activation, making it more difficult to remove adducts^6^, which proportionally affect high-mass ions more^7^. Advances in this area have been made through several approaches, for example modifying the ion, implementing surface induced dissociation (SID)^8^ or combining CID with ion. Lastly, these assemblies are often constructed from varying ratios of subcomponents, each of which can be subject to numerous protein modifications. The resulting *m/z* spectra commonly display broad, unresolved distributions, that are the result of overlapping charge state peaks, precluding mass determination via traditional means. Sometimes subjecting such complexes to tandem MS can disentangle overlapping distributions^9,10^.

One approach to deciphering heterogenous, high mass samples is through single-ion measurements where both *m/z* and *z* (i.e. charge) of an individual ion can be determined: Charge Detection MS (CDMS). Here, the charge of single ions is determined non-destructively using inductive detection. The earliest and simplest application of this principle is single-pass CDMS, first demonstrated in 1960 in order to investigate the collisions of particles with satellites^11^. The term CDMS was coined in 1995, when these principles were combined with electrospray ionization (ESI) and applied to biomolecules, namely DNA of 1-100 MDa^12^. Although, single-pass CDMS is very fast, its charge accuracy is generally limited due to background noise^13^. One way to combat this is to perform multiple measurements on one ion, as the uncertainty is proportional to the inverse square root of the number of measurements. A linear array CDMS setup accomplishes this by chaining multiple detector cylinders, with each detector measuring the charge thus decreasing uncertainty^14,15^. Repeated measurements can also be accomplished by using ion trapping CDMS. In an electrostatic linear ion trap (ELIT) ion oscillations induce charge on a detection cylinder located between two ion mirrors^16,17^. Time-domain data is Fourier transformed (FT) into the frequency domain. Here, the *m/z* of an ion can be inferred from the fundamental oscillation frequency, while the magnitude of said signal is proportional to its charge. The recent ELIT version has reached the charge root mean square deviation (RMSD) of ∼0.2 *e* at ∼1.5 s trapping time, allowing for perfect charge determination^18–20^. A similar ELIT device can be found in the recently commercialized Xevo™ CDMS instrument (Waters Corporation), shown to operate with *m/z* resolution of ∼200, which is appropriate for large biomolecular ions, where mass heterogeneity usually dominates^21^. The most recently constructed ELITs have further improved *m/z* resolution up to 14,600, expanding ELIT application space even further^22^. Other ion traps based on inductive detection can also be adapted to perform simultaneous *m/z* and *z* measurements on single ions, as demonstrated by Orbitrap™-based CDMS. Here, ions are trapped radially around a central spindle electrode and oscillate axially along its axis. Two equal-length, coaxial, quasi-cylindrical inductive detection elements surround the spindle and detect the axial oscillations, allowing for high resolution *m/z* measurements following FT. Though originally constructed for high resolution *m/z* measurements of trapped ion packets, the Orbitrap also allows for charge determination of individual ions if beam intensity is controlled appropriately^23,24^. During trapped-ion CDMS analyses, ions may experience collisions with background gas and losses of neutral molecules (e.g. solvent) resulting in energy and oscillation frequency changes. These phenomena can influence both *m*/*z* and *z* measurements. In an ELIT, ion trajectories may become unstable if their energy shifts beyond the stability region of the device, though the charge induced on the detection cylinder remains unaffected by the trajectory. In contrast, within an Orbitrap device, energy losses reduce the radial trapping radius, leading to reduced charge precision. Moreover, signal loss from ion losses, or changes in ion’s *m*/*z* may occur during prolonged trapping. To resolve this problem and track individual ions in Orbitrap-based CDMS, Selective Temporal Overview of Resonant Ions (STORI) method was developed. Here, the integrated charge is plotted over the detection time at the specific frequency of the ion in question. The resulting slope is proportional to its charge^23^. The STORI method was commercialized through the *Direct Mass Technology* (DMT™, ThermoFisher Scientific)^25^. At 1.5 s trapping time, the expected charge RMSD is ∼1.7 *e*. Using custom data acquisition systems that enable ultra long trapping times, a charge RMSD of 0.55 *e* after 25 s of trapping was achieved^26^.

As mentioned before, viruses are one of the premier offenders when observing heterogeneities precluding charge state assignment in traditional nMS analysis. Studying them, or their capsids, yields important information for pharmaceutical companies, such as quality control in vaccine and vector development^27–29^ and fundamental research. As such, they have been subjected to various CDMS investigations. One such virus is the human norovirus (hNoV), the primary cause of viral gastroenteritis globally and cause of ∼200,000 deaths annually^30–32^. hNoV-derived virus-like particles (VLPs) are self-assembling capsid structures, build from the single major capsid protein VP1. hNoVLPs generally adhere to Caspar-Klug theory^33^ with reported *T =* 1, *T =* 3 or *T =* 4 assemblies with 60, 180 or 240 VP1, respectively, and range from 3 to 14 MDa in a strain-dependent manner^34–37^. Other stoichiometries have been observed occasionally^38–40^ though the native virions form *T =* 3 particles^41^. The N-terminus of VP1 contains double or triple methionine (M) in an MMM or MXM pattern. These methionines can be truncated leading to small differences in the dimer mass and ultimately the particle mass. These masses overlap and, together with the mechanisms described before, preclude charge state resolution. hNoVLPs are non-infectious, stable for elongated times at 4°C^42^ and well characterized^34,43^, making them highly interesting for pharmaceutical^44,45^ as well as fundamental research. Here, we utilize these hNoVLPs to compare two functionally distinct methods of mass determination, namely the single-ion Orbitrap-based CDMS with DMT, which we combine with UniDec deconvolution, and the ELIT-based Xevo CDMS instrument.

## Methods

### VLP production

Full-length VP1 genes of hNoV variants GI.1 Norwalk and GII.17 Kawasaki were cloned and expressed in insect cells using the baculovirus system, as described previously^46,47^. For the GI.1 Norwalk variant (GenBank accession number Q83884), one construct also included the VP2 gene (GeneBank accession number Q83885), while the GII.17 Kawasaki 308 variant (GenBank accession number LC037415.1) included only VP1. Recombinant bacmid DNA was transfected into Sf9 insect cells and incubated at 27 °C for 5–7 days. The culture medium was collected and centrifuged at 3000 rpm for 10 min at 4 °C to recover baculovirus-containing supernatant. This supernatant was then used to infect Hi5 insect cells, which were incubated for 5 days at 27 °C. Following infection, the medium was centrifuged sequentially at 3000 rpm for 10 min and at 6500 rpm for 1 h, both at 4 °C. The supernatant was then concentrated by ultracentrifugation at 35,000 rpm for 2 h (Ti45 rotor, Beckman) at 4 °C. The hNoVLPs were further purified using CsCl equilibrium gradient ultracentrifugation at 35,000 rpm for 18 h (SW56 rotor, Beckman) at 4 °C. Finally, the purified particles were pelleted by centrifugation at 40,000 rpm for 2 h (TLA55 rotor, Beckman) at 4 °C and resuspended in PBS (pH 7.4). VP1 monomer weight was verified using sodium dodecyl polyacrylamide gel electrophoresis (SDS-PAGE)^48^. Staining was performed using a solution containing 0.5 % Coomassie brilliant blue R250, 50 % ethanol and 7 % acetic acid.

### Peptide mapping

Peptide identification was performed using 100 pmol of sample, diluted in 1:9 manner using a dilution buffer (40 mM Tris, 150 mM NaCl, pH 7.5) and injected into an online HPLC system (Agilent Infinity 1260, Agilent Technologies, Santa Clara, CA, USA) using the LEAP HDX PAL robot (AXELSEMRAU by Trajan). This includes a home-packed pepsin column (IDEX guard column with 60 µl Protozyme immobilized pepsin beads, Thermo Scientific, Waltham, MA, USA), a peptide trap column (OPTI-TRAP for peptides, Optimize Technologies, Oregon City, OR, USA), and a reversed-phase analytical column (PLRP-S for Biomolecules, Agilent Technologies, Santa Clara, CA, USA). The HPLC system was directly connected to a timsTOF Pro (Bruker Daltonics, Billerica, MA, USA). The digest was stopped after 180 s. Elution from the analytical column took place over a 10 min linear gradient from 95% solvent A (H_2_O, 0.1% formic acid) and 5% solvent B (acetonitrile, 0.5% solvent A) up to 40% solvent B followed by a step gradient to 80% solvent B for 4 min. Peptides were identified using a data dependent acquisition (DDA) MS/MS method with activated trapped ion mobility and analyzed in PEAKS Studio 10.6 (Bioinformatics Solutions Inc.). In addition to VP1 and VP2, the baculovirus proteome (GeneBank accession number L22858) was also searched against.

### Sample preparation for nMS and CDMS

GI. Norwalk and GII.17 Kawasaki VLPs were exchanged into MS-compatible ammonium acetate at 150 mM (99.99% purity, Sigma-Aldrich) and pH 7.5. The exchange was performed by applying two-step centrifugal gel filtration (Bio-Spin® P-6 size-exclusion columns, 6000 MWCO, Bio-Rad). Monomeric concentration was adjusted to 5 µM prior to data acquisition.

### Orbitrap-based CDMS analysis of hNoVLPs

Nano-ESI capillaries were fabricated from borosilicate glass capillaries (outer diameter: 1.2 mm; inner diameter: 0.68 mm; World Precision Instruments) using a micropipette puller (P-1000, Sutter Instruments) equipped with a squared-box filament (2.5 mm × 2.5 mm) in a two-step pulling program. The pulled capillaries were subsequently coated with a thin layer of gold using a sputter coater (EM ACE600, Leica) under the following conditions: 5.0 × 10^−2^ mbar pressure, 30.0 mA current, and 200 ms. Final capillary opening was performed manually under a stereomicroscope using precision tweezers.

Orbitrap-based CDMS experiments were performed on a Thermo Fisher Q Exactive (QE) UHMR utilizing the DMT mode. For hNoVLPs, the instrument was set to positive ion mode and a trap setting of 7, which relates to a UHV (ultra-high vacuum) read-out of approximately 3.2 × 10^−10^ mbar. Capillary voltage was set between 1.35 kV and 2 kV with 250°C capillary temperature. In-source desolvation was turned on and set to -50 V and HCD collisional energy (CE) set to 100 V. ‘Detector optimization’ and ‘ion transfer target m/z’ were set to ‘high m/z’. The injection flatapole DC, inter flatapole lens, bent flatapole DC, transfer multipole DC and C-trap entrance lens inject were set to 10 V, 10 V, 4 V, 4 V, and 9 V respectively. For single ion acquisition, direct Mass mode was turned on using a resolution setting of 200,000, which equals ∼1 s of trapping time Inject time was manually set to 800 ms. Data acquisition in total spanned 10-15 min, sometimes through several measurements, aiming for >10,000 ions.

### Charge calibration for Orbitrap-based CDMS

To calibrate the relationship between charge and intensity, four well-known samples were chosen: pyruvate kinase (from rabbit muscle, Sigma-Aldrich, P9136-25KU), ß-galactosidase (from *Escherichia coli*, Sigma-Aldrich, G3153-5MG), alcohol dehydrogenase (from *saccharomyces cerevisiae*, ADH, Sigma-Aldrich, A7011-7.5KU) and GroEL (from *Escherichia coli*, Sigma-Aldrich, C7688-1MG). GroEL is approximately 13 times smaller than the capsids anlyzed here. Single ion spectra were recorded for each calibrant and the sum of their spectra was evaluated by UniDec. The mass, minimum and maximum charges were put into the calibration tool of UniDec Charge detection (UCD). The resulting slope of the intensity-charge-plot (Fig. S1) then functioned as a calibration value used in the analysis.

### Analysis workflow for Orbitrap-based CDMS data

Data acquired on the QE UHMR in the form of .raw files was imported into STORIboard (Proteinaceous) and transformed into .dmt files. This allows merging of different .raw files and thus the addition of multiple measurements in order to end up with an appropriate number of ions. Commonly ∼10,000,000 were collected before the filtering steps within STORIboard, where 70 to 80% were removed due to not achieving minimum duration in the Orbitrap. Further filtering steps reduced this number to 10,000 to 30,000.. The processing template for ions greater than 500 kDa was modified, allowing for frequency correction and the number of averaged ions was set to 1. Calibration was set to the default template included in STORIboard, which applies a calibration coefficient of 50,000.

The resulting .dmt files were loaded into the CDMS module of UniDec (UCD)^49^.Charge calibration was first applied, followed by Gaussian smoothing of the *m*/*z* values at 10 and charge values at 1. The resulting 2D charge/mass histograms were binned at 60 kDa and 1 z. 60 kDa mass binning was selected based on VP1 mass. Parameters for the UCD deconvolution algorithm were set to 20 Th for the *‘m*/*z* Spread FWHM’ and 15 charges for the ‘Charge Spread FWHM’ settings. These values were extrapolated from Kostelic et al. and their analysis of AAVs using UCD^49^. Charge state smoothing in the mass dimension and artifact suppression were disabled.

### Xevo CDMS analysis of hNoVLPs

A charge detection mass spectrometer (CDMS; Waters Corporation, Wilmslow, UK), equipped with an electrostatic linear ion trap was used for the experiments. Samples were injected into a nano-ESI cartridge (emitter inner diameter: 5 µm) and ions were generated in positive ion mode. Ions were trapped for 100 ms and the detected time-domain signals are Fourier transformed; the measured frequency and the magnitude correspond to an individual ion’s *m*/*z* and *z* respectively, enabling direct calculation of mass values (*m*/*z * z*). Detected ion signals are filtered in real time. The filter removes ions that were not trapped for the duration of the entire trapping event, ions with an incorrect harmonic ratio (noise peaks) and ions whose frequency shifted beyond the set limit. Filtered data are displayed and updated in real time. Histograms of mass, *m/z* and *z* are updated and visualized in real time using the provided CDMS Data Viewer application. Between 2,500 and 7,000 ions were detected in total. Approximately 40% passed quality filters and were therefore included in the analysis. Histograms of mass, *m/z* and *z* are updated and visualized in real time using the provided CDMS Data Viewer application. Histogram bin width can be adjusted by the user (here bin of 60 kDa was selected).

### Charge and *m*/*z* calibration for Xevo CDMS

In order to calibrate the instrument, data was acquired for proteins spanning an *m/z* range of 4,800 to 14,200 and a total of 13 charge states from *z*=18 to *z*=66. These proteins were enolase (E6126, from baker′s yeast, Sigma Aldrich), Waters™ mass check mAb standard (186006552, Waters Corporation), ß-galactosidase tetramer and octamer (G5635, from Escherichia coli, Sigma-Aldrich), approximately 10 times smaller than the capsids analyzed here. The calibration procedure for the Xevo CDMS instrument is described in more detail by Richardson et al.^50^ but we summarize the main details here.

For *m*/*z* calibration, histogrammed *m/z* spectra are peak detected on an uncalibrated *m/z* axis. The average mass of each protein is assumed to be the sequence mass plus an unknown additional average mass resulting from adducts. The single *m*/*z* calibration parameter is simply a multiplicative rescaling of the provisional *m*/*z* axis, and is determined probabilistically along with an associated uncertainty. Charge is assumed to be related linearly to the amplitude of the fundamental frequency peak for each charge state of each protein. A representative amplitude is determined for each charge state along with an uncertainty and the values obtained from all proteins are used to determine the two parameters of the linear relationship along with associated uncertainties. In the dataset used to calibrate the instrument used in this study, the RMS *m*/*z* calibration residual was 405 ppm and the RMS charge residual was 0.1 *e* (Fig. S2).

### FWHM analysis

Mass spectral data were analyzed using a custom-written Python script. The script processes text files containing two columns: *m*/*z* values and corresponding intensity values. The analysis was restricted to the 8-16 MDa region, where peaks of interest are expected to occur near 10 MDa. Prior to peak analysis, the spectra were smoothed using a Savitzky-Golay filter to reduce noise while preserving peak shape. Within the specified mass range, the script identifies the most prominent peak and extracts three key parameters: the precise *m/z* position of the peak maximum, the peak height (intensity), and the full width at half maximum (FWHM). The FWHM is determined by locating the points on either side of the peak at which the signal reaches half of the maximum intensity. Because the observed peaks are non-Gaussian, the distances from the peak maximum to the half-maximum points on the left and right sides are calculated independently and summed to obtain the total FWHM. Finally, summary statistics including mean values and standard deviations/standard errors of peak positions and FWHM measurements are computed across all analyzed spectra.

### Collection of Cryo-EM micrographs

For cryo-EM measurements, 3.5 µL of sample (VP1 10 µM) was applied to a glow-discharged Quantifoil 300 mesh, R1.2/1.3 copper grid with an extra 2.4 nm graphene layer. Sample was incubated for 5 s before excess liquid was blotted for 3.5 s and the grid was plunged into liquid ethane using the Thermo Fisher Scientific Vitrobot Mark IV system at 8°C and 100% humidity. hNoVLPs were transferred to a 300 kV Titan Krios transmission electron microscope (Thermo Fisher) equipped with an X-FEG electron source and a K3 electron detector camera (Gatan) and a Bioquantum post-column energy filter operated in zero-loss imaging mode with a 20 eV slit width. Data was collected at a pixel size of 0.8 Å with a total dose of 51 *e*^*-*^/Å^2^ using SerialEM^51^. Motion correction, contrast transfer function (CTF) estimation, particle picking and 2D classification was performed in CryoSPARC^52^.

## Results and Discussion

### A practical CDMS workflow strategy ensures comparability

To ensure accurate and reproducible CDMS measurements across two platforms, we established a standardized workflow. hNoVLPs were produced in insect cell culture and purified, as has been demonstrated in previous publications^40,53^ (Fig. 1A). This is followed by quality control using SDS-PAGE, where VP1 from hNoV can be found between 50 and 60 kDa, and negative stain electron microscopy (EM) to determine capsid integrity (Fig. 1B). Additionally, VLP preparations are subjected to bottom-up MS to verify the SDS PAGE and probe for contaminations. The purified sample is buffer-exchanged into a buffer surrogate compatible with nMS, here ammonium acetate. The sample is then diluted to a working concentration of 5 µM (Fig. 1C). Transfer into the gas phase and introduction into the instruments was performed via nano-ESI with either an in-house produced gold-coated capillary or via syringe and the Waters nano-ESI cartridge. (Fig. 1D). Data acquisition on the QE UHMR was performed with Tune. The resulting .raw files were processed using STORIboard, transforming them into .dmt files. These were imported into UCD for deconvolution. The Waters CDMS application software for the Xevo CDMS instrument supplies the user with live data acquisition and analysis (Fig. 1E). In order to determine the true mass of the samples, both systems require calibration (Fig. 1F). For UCD, three or more samples with known masses and charge state resolved spectra are needed to create the calibration curve, relating charge and intensity (Fig. S1). For the Xevo CDMS instrument, *m/z* and *z* calibration is carried out using three protein standards (Fig. S2). Both software applications allow review of spectra and scatterplots for mass, charge and *m/z* (Fig. 1G).

**Figure 1:**
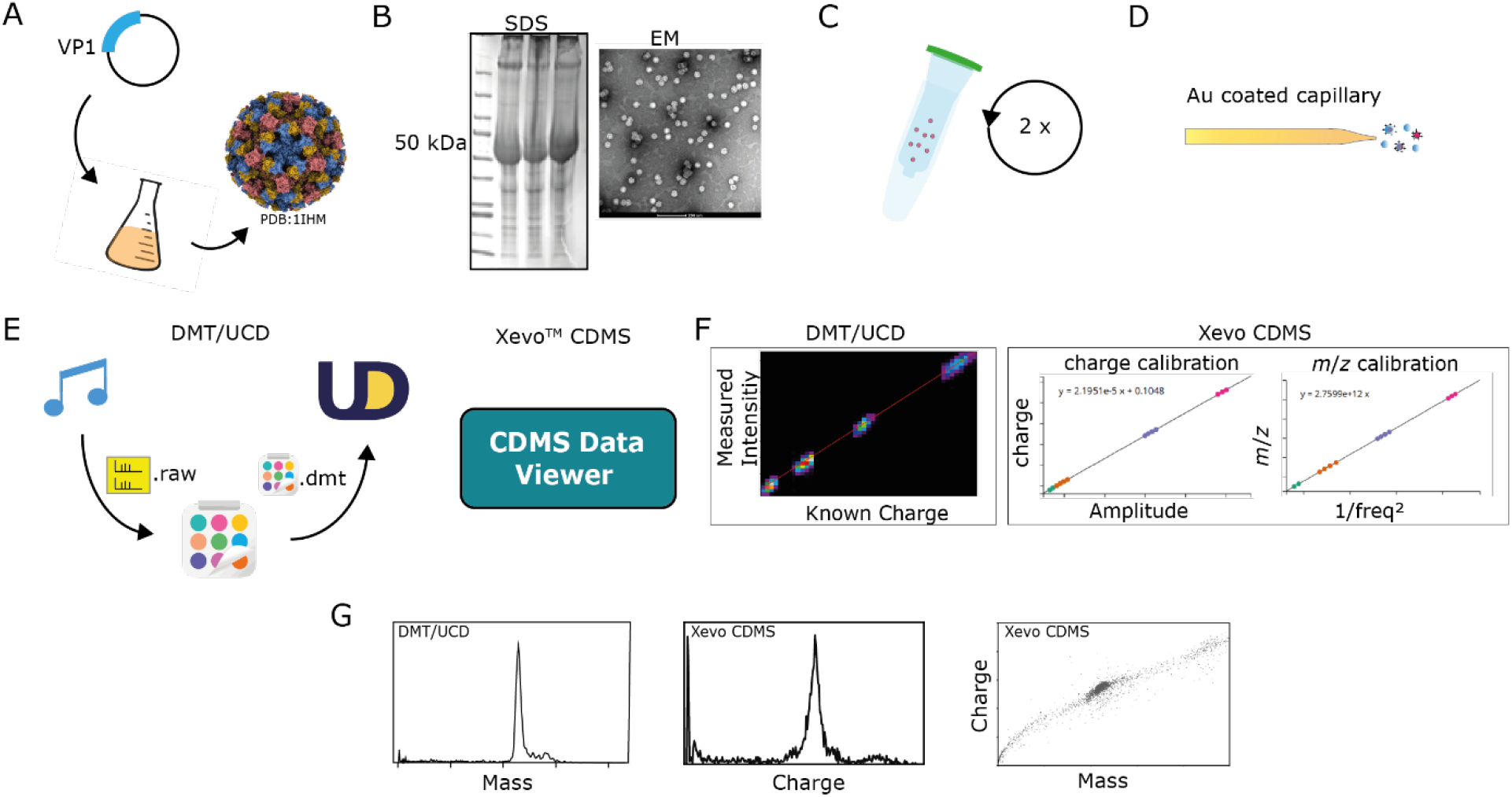
CDMS workflow for hNoVLPs. A) A baculovirus vector containing VP1 (and VP2) is transfected into insect cell culture, from which the self-assembling VP1 is expressed and capsids purified. B) Quality of purification is asserted by SDS-PAGE and EM, shown is GI. 1 Norwalk. C) Buffer exchange into ammonium acetate is achieved by a two-step protocol using spin columns. D) Sample is introduced into the instrument using nano-ESI with either gold-plated, in-house fabricated borosilicate capillaries for the QE UHMR or via syringe injection into the Waters nano-ESI cartridge for the Xevo CDMS^47^. E) Data analysis for DMT mode requires the collected data in Tune to be transformed into .dmt using STORIboard from Proteinaceous. When using UniDec, this file can then be imported into the CDMS module UCD. The CDMS Data Viewer software can be used to control and view data from the Xevo CDMS ELIT instrument. F) Intensity to charge calibration for DMT is performed in UCD by using samples with well-established sizes and charge state resolution. For the Xevo CDMS, both *m*/*z* and charge were calibrated. G) Besides the *m*/*z* spectrum, both of these workflows yield mass and charge histograms and 2D-scatter or density plots combining either of the resulting ion properties.

### Mass determination reveals classical and non-classical assemblies

Both instruments show a major mass distribution for GI.1 Norwalk and GII.17 Kawasaki around 11 and 10 MDa, respectively (Fig. 2A & B). These most likely correspond to *T* = 3 assemblies that can be discerned from negative stain EM (Fig.1, Fig. S7). The masses around 0.12-0.15 MDa match closely with the expected dimer masses for the two strains of 0.11 MDa for GI.1 Norwalk and 0.12 MDa for GII.17 Kawasaki based on the monomer sequence mass (Tab. 1). The slight excess mass here may be caused by the high-mass optimized settings, especially in the DMT method, such as trapping pressure and ion optics. The dimer mass was confirmed by conventional nMS (Fig. S3). GII.17 Kawasaki main peak is 10.92 ± 0.05 MDa as measured by DMT, exceeding the theoretical mass of a *T* = 3 geometry at 10.68 MDa by 2.25%. A similar delta of 2.87% is found in the putative *T* = 4 at 14.66 ± 0.06 MDa compared to the sequence mass of 14.25 MDa, suggesting a systematic overestimation from the analysis. Part of the overestimation is likely due to counterions and incomplete desolvation, usually accounting for ∼1% according to literature^7^. The additional excess mass may stem from the charge calibration that lacks calibrants in the 10 MDa mass range. Charge calibration for DMT requires charge state resolved analytes, which exist but are usually not commercially available. Interestingly, the masses of the Xevo CDMS instrument are smaller than those from DMT. The Xevo CDMS instrument determines the *T* = 3 at 10.64 ± 0.01 MDa which is 0.3% lower than the theoretical mass. The mass accuracy on a calibrated Xevo CDMS is expected to be within ±1%^50^ allowing for the deviation above. In contrast the DMT determined mass is ∼2.6% higher. The difference may be due to the calibration procedures employed and warrants future investigation. Since both samples originated from the same expression, it can be assumed that both adhere to the same stoichiometry of a *T* = 3 assembly. Moreover, conventional nMS data (Fig. S3) excludes N-terminal truncation as explanation for the mass deviation. A *T* = 3 capsid as the main species is in agreement with conventional nMS^39,40^, atomic force microscopy data^54^ and our own EM data (Fig. S7). Besides the aforementioned capsids, GII.17 Kawasaki displays several mass distributions, which do not adhere to the Caspar-Klug principles that govern viral capsid assembly^33^. These mass distributions can be found at 12.98 ± 0.03 MDa / 12.58 ± 0.02 MDa and 13.72 ± 0.07 MDa / 13.32 ±0.01 MDa for DMT/ Xevo CDMS, respectively. They likely represent non-icosahedral capsids, which are either composed of excess or sub-stoichiometric VP1 dimers, leading to tube-like structures with capped ends or incomplete capsids^38^. The Xevo CDMS instrument additionally detects a mass at 20.37 ± 0.04 MDa. This mass does not correspond to any icosahedral symmetry and may either be a non-icosahedral capsid, an aggregate of partially assembled capsids or even a *T* = 3 dimer, although the mass deviation is in favor of the first two options. The main peak of GI. 1 Norwalk at 11.34 ± 0.05 MDa in DMT and 11.04 ± 0.03 MDa in Xevo CDMS displays a large discrepancy to the theoretical mass at 10.18 MDa. This can be observed in both instruments, ranging between 8 and 10% and far exceeding the observed difference in GII.17 Kawasaki (Fig. 2A & B, red). While masses measured using the Xevo CDMS instrument are lower, the offset from the expected mass of ∼1 MDa is comparable in Xevo CDMS and DMT. The EM data (Fig. 1, Fig. S7) however suggest mainly *T* = 3 particles. The main difference between the expression constructs is the presence of the minor capsid protein VP2 in GI.1 Norwalk. VP2 is the second and minor structural protein in noroviruses, and was co-expressed with VP1 previously, while being undetected in structural methods^55^. This may be due to low copy numbers^56,57^, though the interaction with VP1 has been demonstrated multiple times^56,58,59^. It is also hypothesized to form a portal for infection purposes, though this has only been shown in the virions of feline calicivirus (FCV)^60^. To probe if this size deviation could be attributed to VP2, bottom-up MS was applied to both strains. While the sequence coverage for VP1 was 100% in both, an additional 47% coverage for VP2 was found in GI.1 Norwalk (Fig. S4), which was not detected in GII.17 Kawasaki in line with its absence from the expression construct. As VP2 is 22 kDa^61^, our data would suggest ∼36 VP2 become incorporated into the GI.1 VLPs, which would be far above the suggested 1 to 8 copies^56,57^. The discrepancy could be explained with the presence of another protein. Therefore, a plasmid containing only VP1 was generated, omitting the previously included VP2. DMT on this capsid did not yield any sizable difference in mass compared to the original construct (Fig. S5), demonstrating that VP2 incorporation is likely not responsible for the mass overshoot. From the baculovirus expression system, there are several proteins detected in bottom-up MS, amongst others glycoprotein 64 (GP64, Fig. S3). Normally, GP64 is attached to the baculovirus envelope^62,63^ but can also be observed in the conventional nMS in the dimer region of GI. 1 Norwalk at 64 kDa (Fig. S3). VP2 and GP64, incorporated either by binding to VP1 or within the hollow shell, may account for at least some of the 10%, or >1 MDa, difference between the expected and determined mass. Though, so far there has been no other evidence for the incorporation of these proteins into the capsids.

**Figure 2:**
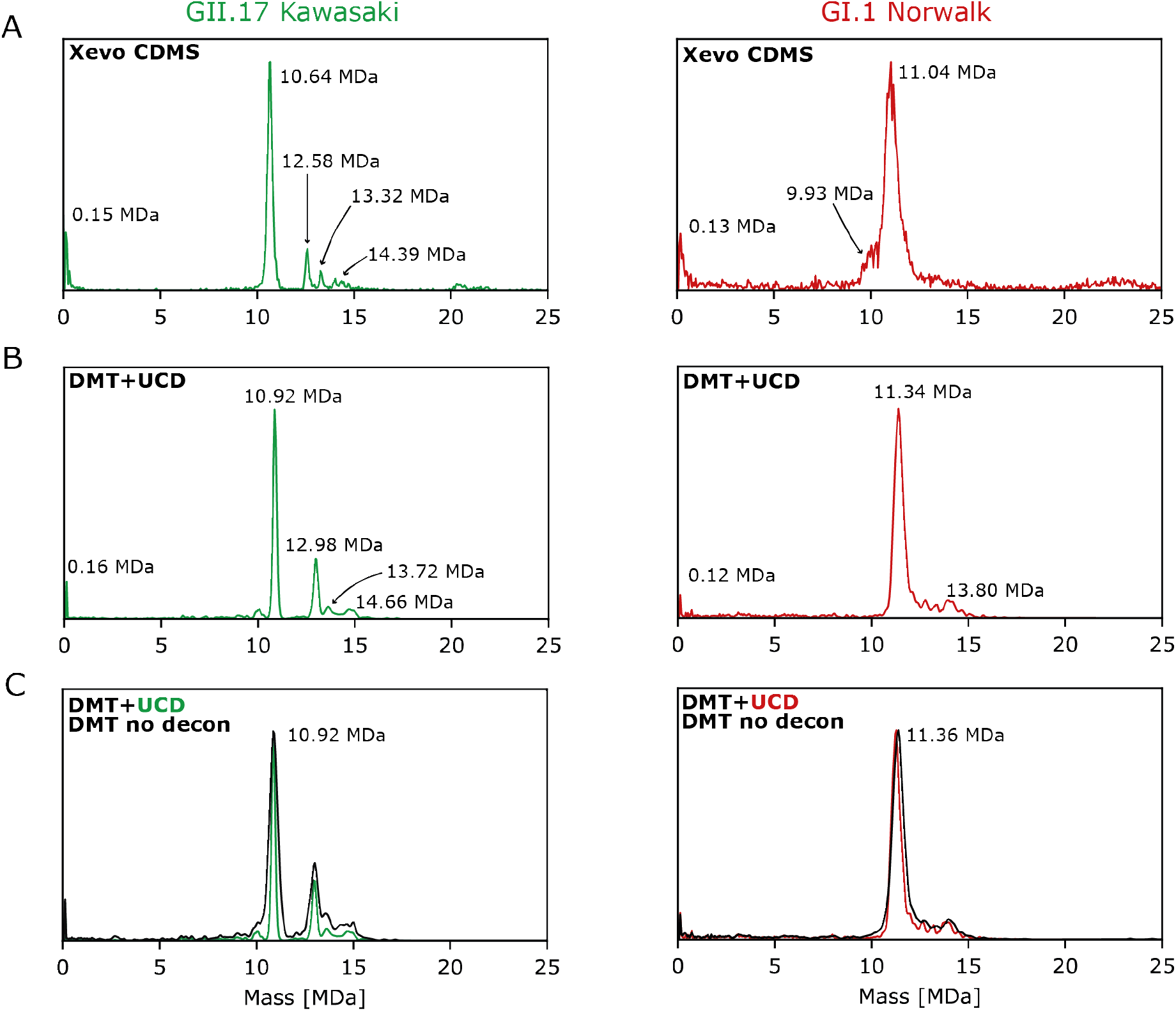
GII.17 Kawasaki and GI.1 Norwalk hNoVLPs in Xevo CDMS and DMT+UCD. Representative mass spectra of hNoVLPs from GII.17 Kawasaki (green) and GI.1 Norwalk (red) in the **A)** Xevo CDMS and **B)** DMT combined with Unidec CDMS deconvolution (UCD). The respective instrument is annotated in each spectrum. Both strains are displaying similar mass distributions in each instrument, though the Xevo CDMS masses are smaller than those detected in the UHMR and are as such closer to the theoretical mass of the detected assemblies. **C)** Comparing the effect of deconvolution on the DMT spectrum. Non-deconvolved spectrum is shown in black with non-deconvolved mass annotated. All annotated masses are means (*n* = 3).

Another capsid can be detected at 13.80 MDa / 13.63 ± 0.28 MDa, DMT/ Xevo CDMS respectively, which are in line with the theoretical mass of *T* = 4 particles. This symmetry has been reported for GI.1 Norwalk^38^, though the signal in the Xevo CDMS instrument is low. If these are indeed *T* = 4, they would be lacking the mass overshoot that can be observed in the putative *T* = 3. In DMT, there are seemingly additional peaks between 11.04 and 13.80 MDa in the deconvoluted data. Comparing these peaks to the non-deconvolved data, it is much less pronounced among the shoulder of the main peak and may be part of multiple non-classical assemblies adjacent to the main peak. Another explanation is that the UCD deconvolution strongly overemphasizes said shoulder peak, creating an artifact signal. Notably, the Xevo CDMS instrument resolves a feature at 9.93 ± 0.03 MDa that cannot be found in DMT. This distribution is 2.46 % smaller than the theoretical *T* = 3 mass and therefore closer than the main peak mass, though the margin of mass underestimation is more suggestive of another prolate rather than the actual *T* = 3 capsid. The Xevo CDMS instrument also measures a mass at 22.61 ± 0.47 MDa (Fig. 2B), close to a *T* = 7 capsid (Tab. 1). The existence of a GI.1 *T* = 7 capsid has been noted before^38^, but the mass here is smaller than expected for *T* = 7 and may belong to an aggregate of two *T* = 3 capsids. This signal cannot be found in the DMT, which may be due to poorer high mass transmission or more aggressive desolvation. It is similar to the putative aggregate found in the GII.17 Kawasaki data.

**Table 1:** Detected masses from both strains and instruments. Table containing all masses detected and theoretical (th) masses for several, non-truncated assemblies for comparison. Data is given as a mean (*n* ≥ 2) with standard deviation. If no deviation is given only one mass was obtained, a deviation of 0.00 denotes no deviation within the data set. The masses are overall similar with most assemblies being detected in both instruments. The masses from the Xevo CDMS are smaller than those from DMT and are closer to the theoretical masses.

|  | GII.17 Kawasaki (MDa) |  |  | GI.1 Norwalk (MDa) |  |  |
| --- | --- | --- | --- | --- | --- | --- |
|  | th. mass | DMT | Xevo CDMS | th. mass | DMT | Xevo CDMS |
| VP1 | 0.05938 | - | - | 0.05658 | - | - |
| VP1<sub>2</sub> | 0.11876 | $0.16 \pm 0.03$ | $0.15 \pm 0.02$ | 0.11316 | 0.12 | 0.13 |
| | | $10.04 \pm 0.06$ | | | | $9.93 \pm 0.13$ |
| $T = 3$<br>(180 VP1) | 10.68 | $10.92 \pm 0.05$ | $10.64 \pm 0.01$ | 10.18 | $11.34 \pm 0.05$ | $11.04 \pm 0.03$ |
| | | $12.98 \pm 0.03$ | $12.58 \pm 0.02$ | | $13.05 \pm 0.15$ | |
| | | $13.72 \pm 0.07$ | $13.32 \pm 0.01$ | | | |
| $T = 4$<br>(240 VP1) | 14.25 | $14.66 \pm 0.06$ | $14.39 \pm 0.04$ | 13.58 | $13.80 \pm 0.00$ | $13.63 \pm 0.28$ |
| | | | $20.37 \pm 0.04$ | | | $22.61 \pm 0.47$ |
| $T = 7$<br>(420 VP1) | 24.94 | - | | 23.76 | | |

### Impact of post-processing

Data processing through the UniDec UCD module drastically reduces FWHM with minimal impact on mass determination (Fig. 2C). Here, GII.17 Kawasaki *T* = 3 presents at 10.92 ± 0.05 MDa and GI.1 Norwalk at 11.36 ± 0.06 MDa. As such, the FWHM of the *T* = 3 peak is decreased from 0.40 ± 0.24 MDa to 0.37 ± 0.05 MDa and from 0.71 ± 0.05 MDa to 0.47 ± 0.08 MDa, for GII.17 and GI.1, respectively (Tab. S1). The deconvolution algorithm compares the measured data to deconvolved data, which at the starting point equals the measured data, and iterates through multiple fits of a point spread function. The deconvolved data updates with every iteration until either a user defined threshold is met or the change in the deconvolved data is so small that convergence is indicated. This virtually reduces charge state uncertainty resulting in increased resolution. The ∼50% decrease in FWHM through deconvolution aligns with the change in FWHM in the original UCD publication^49^. Comparing the FWHM between DMT+UCD and Xevo CDMS (Tab. S2), GII.17 Kawasaki displays similar values over all triplicates of 0.37 ± 0.08 MDa and 0.356 ± 0.004 MDa, respectively. The FWHM for GI.1 Norwalk is 0.47 ± 0.08 MDa in DMT+UCD and 0.83 ± 0.035 MDa in Xevo CDMS, highlighting a much starker contrast. Part of this contrast is due to the deconvolution (FWHM non-deconvoluted data: 0.71 ± 0.05 MDa) but in GII.17 Kawasaki, the FWHM reduction is far less than in GI.1 Norwalk and the deconvolved DMT FWHM is much closer to the Xevo CDMS FWHM. The FWHM in the non-deconvolved GI.1 Norwalk is also narrower than the FWHM in the Xevo CDMS data. This suggests that this is not simply a case of forced data reduction through deconvolution but rather a dataset specific effect in GI.1 Norwalk in the Xevo CDMS instrument. This may point to instrument specific influences on the distribution of molecules causing the excess mass on the putative *T* = 3 GI.1 Norwalk capsid, suggesting that these molecules associate to the particle exterior.

### Charging behavior differs in the instruments

Observing the charging behavior in both instruments can lead to a more refined understanding of the differences in mass determination. Using charge to mass scatter plots (Fig. 3), the overall patterns of both strains present as almost identical in both instruments. They show lower amount of signal in the lowest mass region, corresponding to VP1 dimers, from where scattered data points rise in a curve-like manner to the ∼10 MDa region, which contains most data points. When going further up in mass, the amount of signal lessens again. Around ∼20 MDa, the Xevo CDMS data shows signals assigned to VLP aggregates while the DMT+UCD does not, a difference that was highlighted above as well. GII.17 Kawasaki (Fig. 3A) has the same discernible assemblies in both instruments, with the signal that represents the *T* = 3 capsid forming a tear shape pointing downwards, showing that some ion species here have a lower charge than the bulk of the ions. Interestingly, this seems to be specific to GII.17 Kawasaki and cannot be found in GI.1 Norwalk (Fig. 3B). GI.1 Norwalk charges overall less then GII.17 Kawasaki *T* = 3, which is similar to charging of the streak in the tear shaped distribution. As charging corresponds to displayed surface area^64^, this suggests an enlarged population of the capsid in GII.17 Kawasaki giving rise to the high charge states. The GII.17 Kawasaki *T* = 3 capsid distribution in the Xevo CDMS carries decidedly less charges than 300, while the DMT+UCD capsid has clearly more. A similar pattern can be seen in GI.1 Norwalk, though somewhat less dramatic. A straightforward explanation for this is a difference in charge calibration caused by using smaller calibrants that are too small compared to the analytes used here, causing a deviation in the calibration coefficient, a critical value for proper mass assignment. However, this can only explain small differences as the determined masses are very close to each other. The biggest calibrant in DMT+UCD here is the GroEL 14mer at ∼800 kDa, while Xevo CDMS uses the β-galactosidase octamer around ∼1 MDa. This slightly heavier calibrant could account for this difference, though both have similar charges around 66. Another reason for charge and thus mass overestimation may be the use of nitrogen as a collision gas in the QE UHMR instead of the recommended xenon for heavier ions^65^. Collisions with relatively light gas molecules lead to less effective desolvation and potentially insufficient cooling, thus impacting ion trajectory compared to calibrants and consequently mass determination.

**Figure 3:**
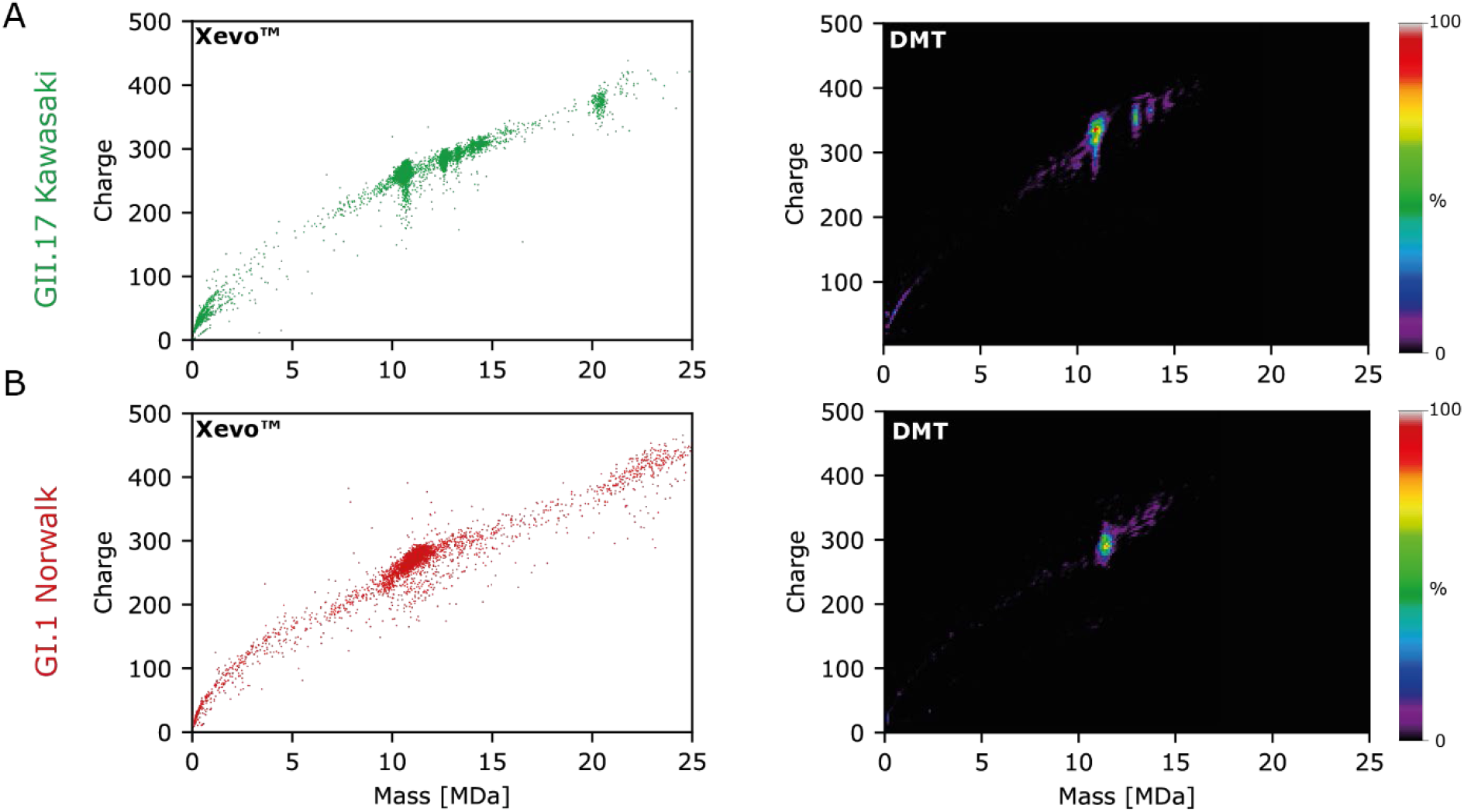
Charge to Mass Plots. Charge to mass scatter plots from the Xevo™ CDMS and 2D-density plots for DMT from UniDec, where the intensity is displayed by the color scheme, for **A)** GII.17 Kawasaki and **B)** GI.1 Norwalk. Charges are slightly lower in the Xevo CDMS compared to the DMT charges. Overall pattern is similar again, with most abundant mass distributions clearly visible and distinguishable in both instruments. Charges in the Xevo CDMS instrument are lower overall, which is especially noticeable in the GII.17 Kawasaki.

A direct comparison showed no mass differences due to collision gas in the instrument employed here (Fig. S6). Another and hence most likely explanation is that the QE UHMR produces higher charge states than other instruments. This has been observed for alcohol dehydrogenase previously, where the main charge state was shifted from 24+ to 25+ reproducibly^66^.

### Sufficient heat reduces mass in Xevo CDMS

The inlet situated behind the entrance of the Xevo CDMS instrument is equipped with heating capabilities, though the initial measurements were performed at room temperature (RT). The distance between heating inlet and capillary is likely too large for the heating to direct impact the capsids in-solution. In order to gain detailed insight into the impact of heat on the assemblies, temperatures were raised in 20°C steps between 100°C and 300°C (Fig. 4A). The mass of GII.17 Kawasaki *T* = 3 rises initially at 100°C to 10.71 ± 0.01 MDa, stays mostly at this mass until it starts decreasing at 240°C until the maximum temperature of 300°C, where it stops at 10.55 ± 0.02 MDa (Fig. 4C, Tab. 2). The prolate capsid at 12.58 ± 0.02 MDa at RT behaves similarly, initially increasing to 12.72 ± 0.02 MDa and ultimately ending at 12.49 ± 0.02 MDa. The mass difference from RT to 100°C is under 1% and may only be inefficient desolvation, although heating should rather aid desolvation and thus improve performance^67^. The low mass region shows a slight loss of signal intensity going from RT to 100°C, which then rises again at 240°C, though not to the original intensity, and then stays at this level. The initial loss of signal intensity could point to a loss of free VP1 dimer, which is being incorporated into or aggregate with the capsid, accounting for the ∼70 kDa extra mass. At a heat threshold, here 240°C, the oversized capsid could be destabilized as disassembly of viral capsids through heat is well documented^68^, and releases the additional VP1 dimer and thus increasing the low mass intensity again. The mass loss beyond the theoretical mass may be due to further subunit loss, though the low signal intensity in the low mass region does not fit this hypothesis.

**Table 2:** Assembly masses throughout heating process. Masses for two assemblies per strain throughout all heating steps. When a standard deviation is given, triplicates were measured, one dataset has been collected for all temperatures. All masses demonstrate an initial rise and then slight loss of mass.

|  | GII.17 Kawasaki (MDa) |  | GI.1 Norwalk (MDa) |  |
| --- | --- | --- | --- | --- |
| | prolate | $T = 3$ | Putative prolate | putative $T = 3$ |
| $100^\circ\text{C}$ | $12.65 \pm 0.01$ | $10.71 \pm 0.01$ | $9.89 \pm 0.18$ | $11.07 \pm 0.04$ |
| $120^\circ\text{C}$ | 12.67 | 10.71 | 9.86 | 11.27 |
| $140^\circ\text{C}$ | 12.68 | 10.73 | 9.62 | 11.11 |
| $160^\circ\text{C}$ | 12.71 | 10.71 | 9.93 | 11.06 |
| $180^\circ\text{C}$ | 12.69 | 10.71 | 10.05 | 10.99 |
| $200^\circ\text{C}$ | $12.72 \pm 0.02$ | $10.73 \pm 0.02$ | $10.18 \pm 0.11$ | $11.05 \pm 0.03$ |
| $220^\circ\text{C}$ | 12.73 | 10.74 | 9.82 | 10.98 |
| $240^\circ\text{C}$ | 12.67 | 10.69 | 10.05 | 10.94 |
| $260^\circ\text{C}$ | 12.63 | 10.64 | 9.88 | 11.01 |
| $280^\circ\text{C}$ | 12.59 | 10.59 | 9.98 | 10.90 |
| $300^\circ\text{C}$ | $12.49 \pm 0.02$ | $10.55 \pm 0.02$ | $9.99 \pm 0.02$ | $10.86 \pm 0.02$ |

**Figure 4:**
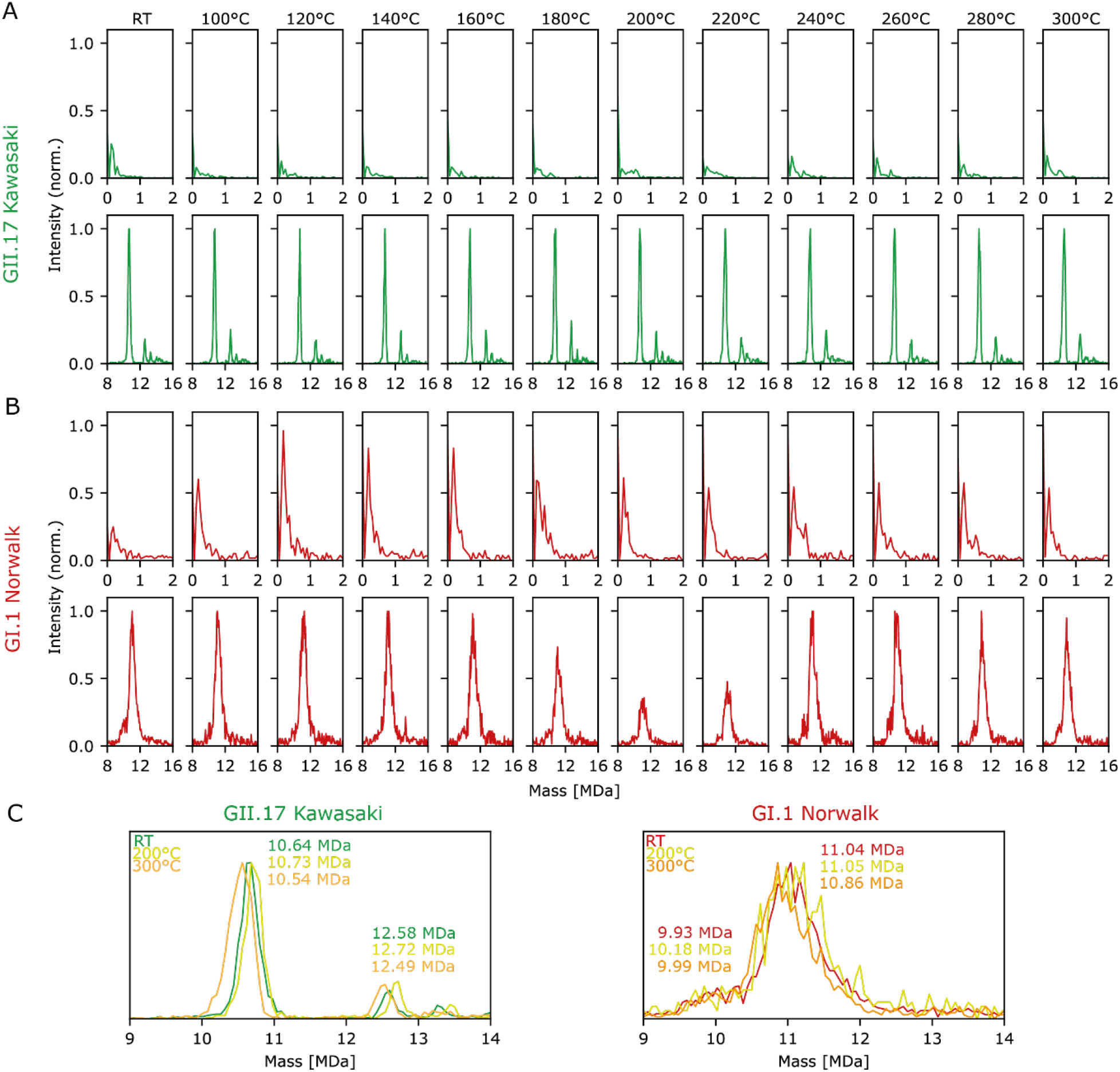
Stepwise inlet heating in the Xevo™ CDMS decreases mass. Applying heat in 20°C steps starting with 100°C up until the maximum temperature of 300°C. Room temperature (RT) is displayed first for comparison. Low mass region, 0 to 2 MDa, and high mass region, 8 to 16 MDa, are shown for **A)** GII.17 Kawasaki and **B)** GI.1 Norwalk. In GI.1 Norwalk heating causes immediate increase in intensity for dimer mass throughout the heating process. Kawasaki low mass spectra do not show a large change. **C)** Overlay of representative RT, 200°C and 300°C spectra with annotated masses.

The overarching pattern is the same in GI.1 Norwalk, with a slight increase in mass to 11.07 ± 0.04 MDa at 100°C, 11.05 ± 0.03 MDa at 200°C and 10.86 ± 0.02 MDa,constituting a mass loss of ∼2%. This is twice the percentage mass loss of GII.17 Kawasaki. The heat stability differences between the strains utilized here has been previously tested with molecular dynamics (MD) and ion mobility, where the GII.17 Kawasaki VP1 dimer was more resilient to unfolding than GI.1 Norwalk^69^. Additionally, the greater relative loss could also be attributed to the loss of VP2 or GP64, if attached to the capsid structure. The dramatic increase of signal in the low mass region at 100°C and 120°C originates from the 0.16 MDa mass originally detected and points to dimer losses, which would agree with the overall mass loss of 220 kDa, or two VP1 dimers. Notably, the mass close to the main peak, at 9.93 ±0.13 MDa also shows the same pattern of mass increase and loss. At 200°C the mass equals the theoretical mass of the *T* = 3 capsid and is subsequently smaller at 300°C. The mass difference is closer to that for the GII.17 Kawasaki *T* = 3 and prolate,∼100 kDa. Stability may also be higher for these than for the main peak in the GI.1 Norwalk spectrum, precluding greater mass losses. This higher stability together with mass underestimation may identify the 9.93 MDa peak as the actual *T* = 3 mass without any bound proteins, though EM shows highly a homogenous capsid population (Fig. S7). Having the same mass loss, throughout 3 different assemblies and two different strains suggests an overarching mechanism for mass loss or desolvation. Interestingly, the FWHM do increase with heating, though for GII.17 Kawasaki only at 300°C and not at 100°C or 200°C. For GI.1 Norwalk, FWHM is the highest at 200°C and slightly lower at 300°C, suggesting that the process of subunit loss may be ongoing at 200°C and finished at 300°C (Tab. S1).

### Cryo-EM data confirms distinct assemblies in GII.17 Kawasaki

While there are already crystal and cryo-EM structures for GI.1 Norwalk VLPs^38,70^, there are none so far for GII.17 Kawasaki. Therefore, cryo-EM was applied in order to inspect the different assemblies more closely. Figure 5 shows a micrograph and 2D classes of three distinct assemblies in GII.17 Kawasaki. The micrograph displays mostly classical particle shapes that can be attributed to *T* = 3 and substantially less prolate and *T* = 4 capsids (Fig. 5A). This is in line with the number of particles in the different 2D classifications: 211,502 particles for *T* = 3, 19,985 particles for the prolate and 30,675 particles for *T* = 4. While the overrepresentation of the *T* = 3 particles fits the CDMS data well, the relationship between prolate and *T* = 4 seems inverse. The prolate may be underrepresented due to the harsh blotting and freezing process destroying some of these potentially less stable capsids, caused by their non-classical symmetry^33^. Representative 2D classification of the three different assemblies that are observed show the expected icosahedral shape for *T* = 3 and *T* = 4 (Fig. 5B & D). The prolate capsid classification demonstrates an oval shape (Fig. 5C). Using the mass from the CDMS data, the prolate is made up of ∼212 VP1 monomers, which is 32 more than the *T* = 3 and 28 less than *T* = 4. A symmetrically allowed prolate *T* = 3 would contain 210 VP1 dimers with an additional ring of hexamers between the endcaps, creating the elongated, oval shape. This agrees with the 2D classes and a similar shape has been reported before in another CDMS and cryo-EM study for GI.1 Norwalk^38^. Though here, the prolate exceeds the allowed number of dimers by two. This may either be the result of an overshoot or the prolate is an intermediate between *T* = 3 and *T* = 4. Thus, as this shape appears both in gas-phase and on a frozen surface, it is unlikely that it is an artifact of either methodology. It is unclear, whether this is a metastable form, a step in-between *T* = 3 and *T* = 4, or a true pseudo capsid on its own, but the abundance and resistance to heat and cold point to the latter. To our knowledge, this is the first time 2D classes for GII.17 Kawasaki *T* = 3, *T* = 4 and a prolate capsid have been reported. The last one is a major step towards the first full structural reconstruction of a norovirus prolate capsid.

**Figure 5:**
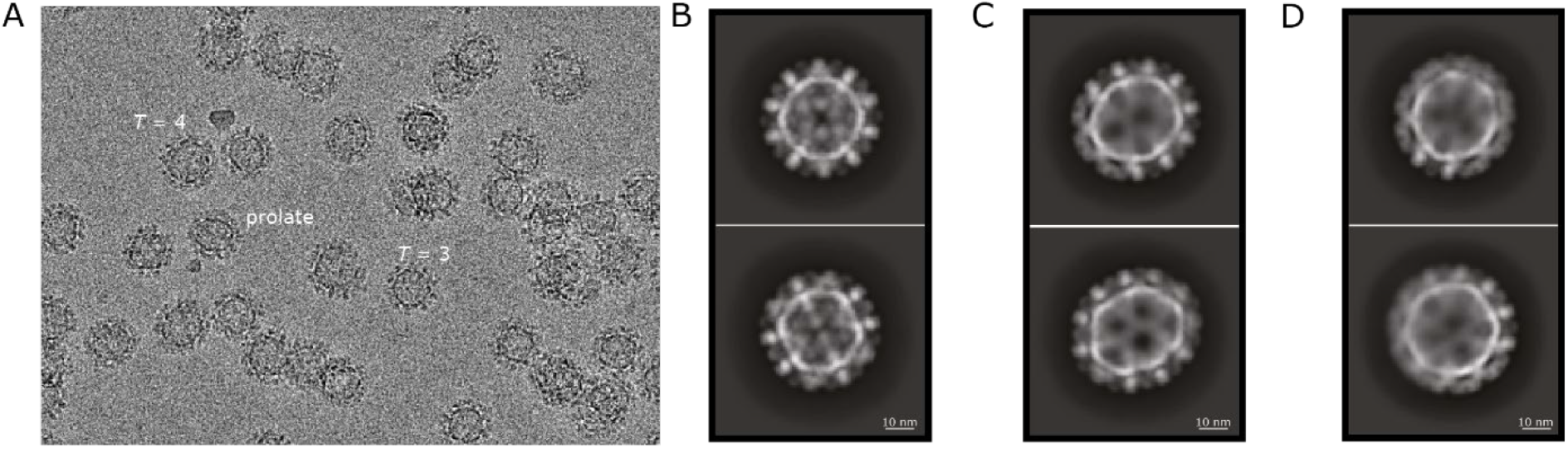
Micrographs and 2D classes of GII.17 Kawasaki VLPs. **A)** Cryo-EM micrograph displaying the three distinct assemblies *T* = 3, the prolate and *T* = 4. **B)** Representative 2D class for *T* = 3 capsid. **C)** Representative 2D class for the prolate capsid, demonstrating the oval shape.**D)** Representative 2D class for *T* = 4 capsid.

## Conclusion

The two CDMS instruments used here identified the same assemblies, though the Xevo CDMS is slightly closer to the theoretical masses expected for hNoVLPs. This could be related to the dual calibration or better desolvation. The DMT coupled with UniDec analysis has a slight edge in apparent mass resolution as a result of deconvolution, but also comes with a more complex workflow. Additionally, deconvolution can result in loss of information about the true width of the mass distribution or in the creation of artificial features and must therefore be applied judiciously. The Xevo CDMS can induce heating within the instrument, affecting the sample in a destabilizing manner, which may be especially useful for stability experiments or assigning peak identity. The mass differences between both workflows likely stem from the *m*/*z* calibration for the Xevo CDMS, which takes into account adduct mass and hence automatically corrects for adducts. Additionally, different charge calibrants can cause deviations in mass, especially in the DMT workflow as the extrapolation to the analyte is slightly higher. Here, the biggest calibrant is approximately 13 times smaller than the capsid while it is ∼10 times smaller for the Xevo CDMS. However, the consistency in mass overestimation in DMT for GII.17 Kawasaki points to a minor contribution to the differences between the systems. Interestingly, the two samples that were tested here behaved somewhat differently than expected. GI.1 Norwalk presented a major mass distribution around 11 MDa that was ∼1 MDa oversized compared to the expected mass at 10.18 MDa. We listed and investigated several hypotheses that all included some sort of protein binding causing the mass overage. The presence of a low intensity mass around 9.9 MDa in Xevo CDMS, close to the expected mass may point to free or non-bound capsid though the mass is slightly too low. We were able to exclude VP2 as a culprit but could not find a definite answer to this, warranting further research. GII.17 Kawasaki on the other hand very clearly assembled into the expected sizes of a ∼10.65 MDa *T* = 3 and ∼14.25 MDa *T* = 4 capsid and additionally an “in-between” sized prolate capsid around 12.58 MDa. These three sizes were then confirmed by cryo-EM, the first of its kind for GII.17 Kawasaki. This work presents an in-depth investigation of two commercially available CDMS technologies with heterogenous, high-mass samples.

## Acknowledgements

The authors would like to acknowledge funding from the EU through H2020 EIC Pathfinder Open Ariadne-Vibe (964553, CU&LT) and HEurope EIC Pathfinder Open, VIRUSong (101099058, CU&RP&SU), from the BMFTR through RÅC SAXFELS 05K22PSA (CU&TD&JCKK) and VirMScan 13GW0622 (CU&JMG), from the Deutsche Forschungsgemeinschaft (DFG, German Research Foundation) through Vision GRK2887 – 497350882 (CU&JR) and through Emerging Viruses SFB 1648/1 (2024)

– 512741711 (CU&WS). CU also acknowledges an equipment grant from the Free and Hanseatic City of Hamburg. The Leibniz Institute for Virology (LIV) is supported by the Free and Hanseatic City of Hamburg and the Federal Ministry of Health (BMG).

## Conflict of interest

Anisha Haris, Jakub Ujma, Keith Richardson and David Eatough are employees of Waters Corporation, who manufactures Xevo™ CDMS instrument. Steve Preece and Kevin Giles are former employees of Waters Corporation.

## Notes

### Competing Interest Statement

Anisha Haris, Jakub Ujma, Keith Richardson and David Eatough are employees of Waters Corporation, who manufactures XevoTM CDMS instrument. Steve Preece and Kevin Giles are former employees of Waters Corporation.

