## Supplemental Material for "Applying distinct CDMS strategies to observe non-classical virus capsid assembly"

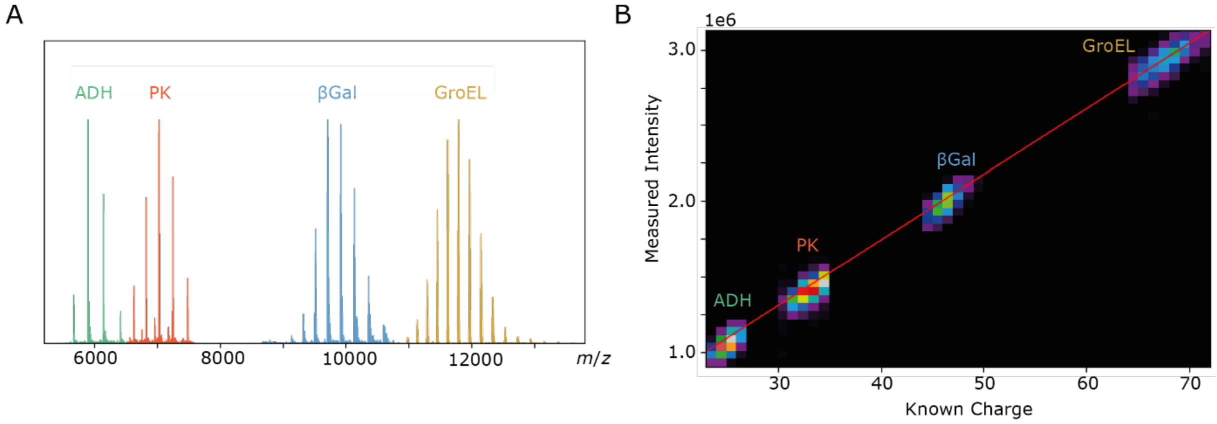


**Figure S1: Charge calibration for DMT. A)** representative nMS of charge calibrants utilized in the charge calibration for DMT in UCD. **B)** Charge calibration relating the known charges of the calibrants with their measured intensities. The slope of the linear regression is the calibration coefficient that is later applied to datasets.

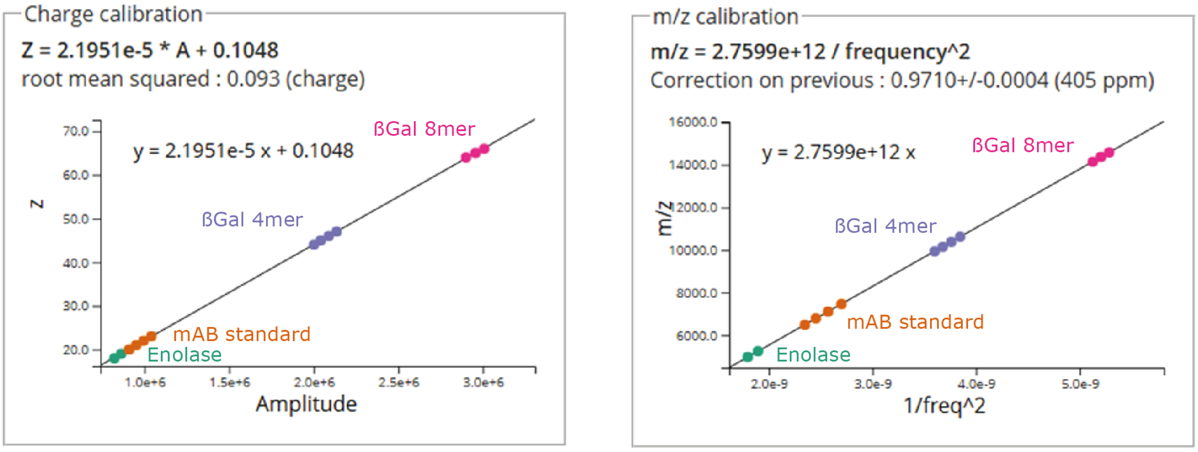


**Figure S2: Charge (left) and *m/z* (right) calibration for Xevo CDMS.** For charge calibration charges of samples, ranging from *z*=18 to *z*=66, were plotted against the respective amplitudes. For *m/z* calibration *m/z* peaks were plotted against inverse squared frequency.

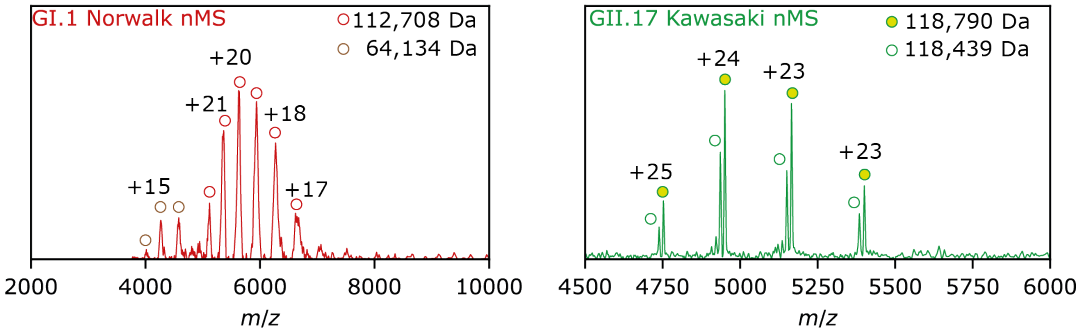


**Figure S3: Native MS of VP1 dimer.** VP1 dimer is measurable in the same sample conditions as full capsids. GI.1 Norwalk has a negative delta of ~500 Da from the theoretical mass, likely due to the truncation of 3 methionines from the N-terminus. The second peak distribution may belong to GP64 from baculovirus used in the expression system. GII.17 Kawasaki shows two very close masses, where the difference is caused by the truncation of the N-terminal methionine and lysine.

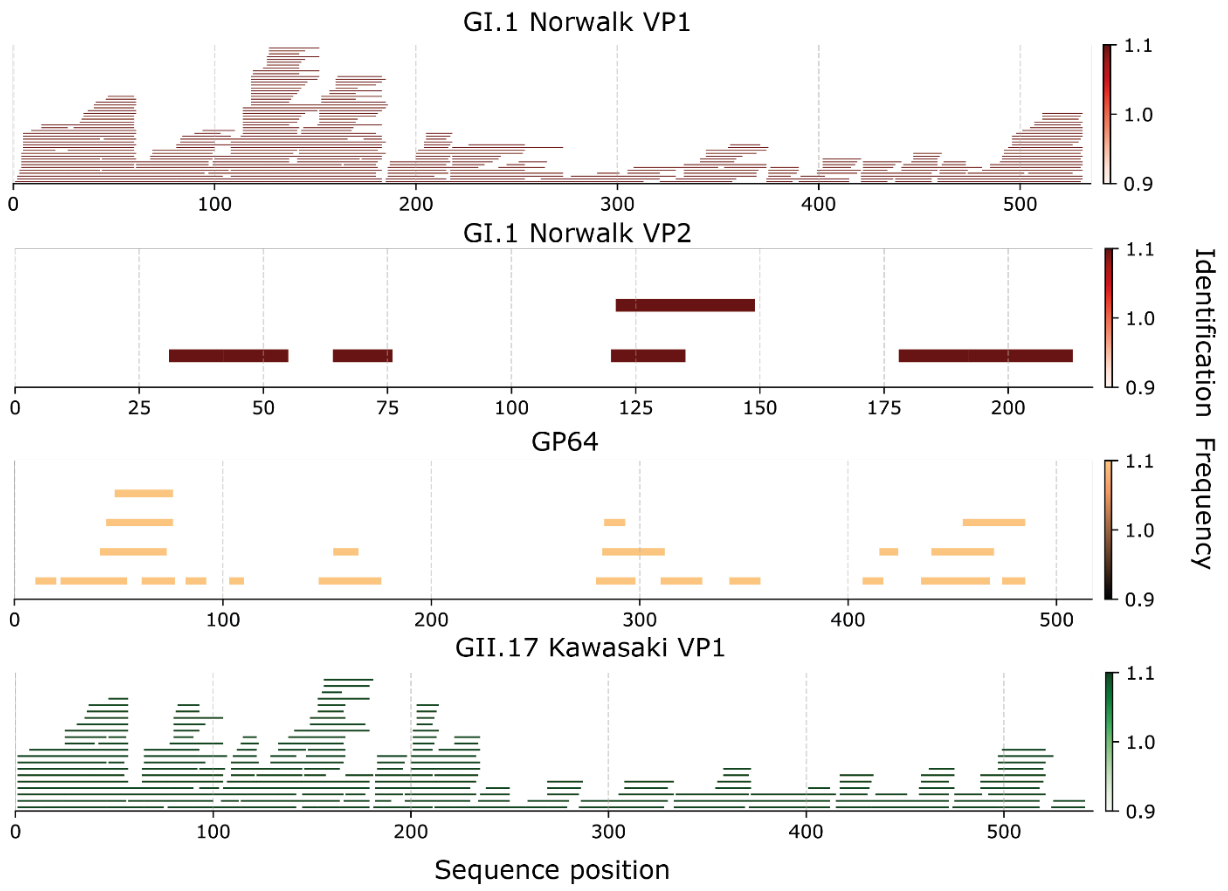


**Figure S4: Peptide mapping of VP1, VP2 and GP64.** Peptide mapping overview from pepsin digestion of GI.1 Norwalk VP1 (100 % coverage), GI.1 Norwalk VP2 (47 % coverage), baculovirus GP64 in GI.1 sample (26 % coverage) and GII.17 Kawasaki VP1 (100 % coverage). Overlapping peptides are stacked.

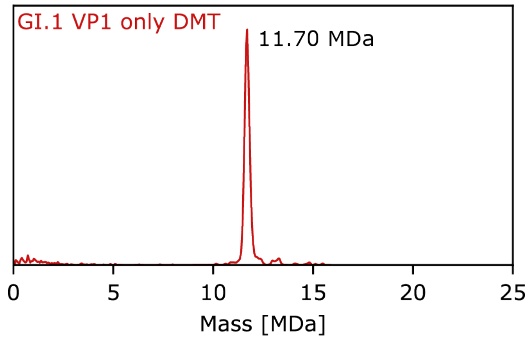


**Figure S5: DMT of GI.1 Norwalk derived from plasmid exclusively encoding VP1.** Expressing a plasmid containing only VP1 does not alter the detected mass of GI.1 Norwalk in DMT, suggesting that VP2 is not the cause for the mass deviation.

**Table S1: FWHM from both instruments.** Full width half maximum (FWHM) as calculated with a custom script. Data is given as a mean (*n* = 3) with standard deviation. GII.17 Kawasaki has an overall smaller FWHM than GI.1 Norwalk and has a similar FWHM on both instruments. GI.1 Norwalk has a much higher FWHM in the Xevo CDMS than in DMT.

| | GII.17 Kawasaki (MDa) | GI.1 Norwalk (MDa) |
|---|---|---|
| DMT non-deconvolved | 0.40 ± 0.24 | 0.71 ± 0.05 |
| DMT+UCD | 0.37 ± 0.08 | 0.47 ± 0.08 |
| Xevo CDMS | 0.356 ± 0.004 | 0.83 ± 0.04 |
| Xevo CDMS 100°C | 0.37 ± 0.03 | 0.833 ± 0.012 |
| Xevo CDMS 200°C | 0.37 ± 0.02 | 0.91 ± 0.06 |
| Xevo CDMS 300°C | 0.43 ± 0.03 | 0.871 ± 0.001 |

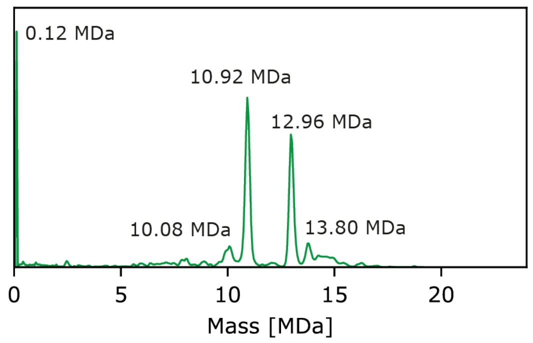


**Figure S6: DMT and UCD of GII.17 Kawasaki using Xe.** Representative mass spectrum showing mass determination of GII.17 Kawasaki with Xe as collision gas. While the overall pattern has changed, most notably more prolate compared to *T* = 3 and more VP1 dimer appear, the masses remain the same.

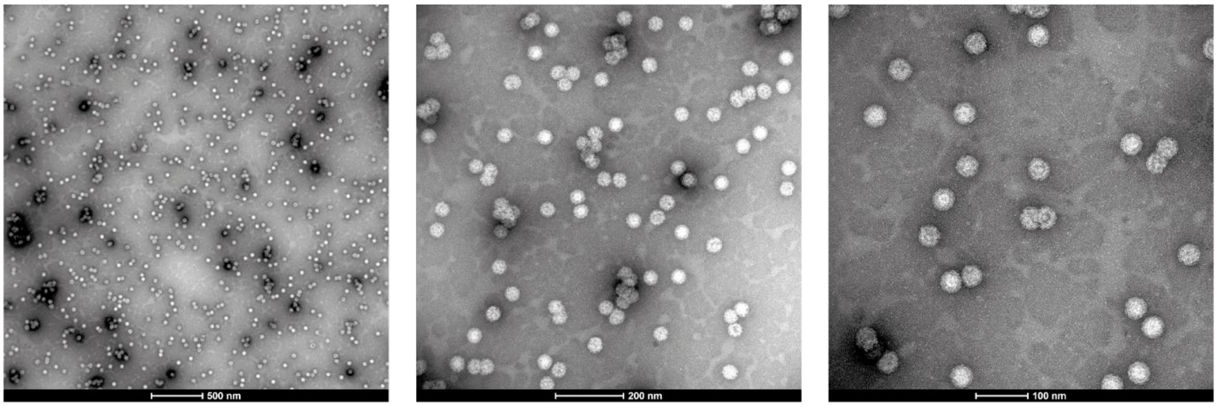


**Figure S7: Negative stain EM at different magnifications.** EM data of GI.1 Norwalk at three different magnifications as indicated in each micrograph. The capsids are mostly homogenous and predominantly *T* = 3.
